# Modeling IK1 current modulation by melatonin and luzindole: a benchmark for patch clamp studies

**DOI:** 10.64898/2026.02.04.703771

**Authors:** Mikhail Anatolyevich Gonotkov, Victoria Faizovna Safiulina

## Abstract

Whole-cell patch-clamp studies often fail to observe the expected effect of melatonin on the IK1 current in cardiomyocytes, which may be due to cytoplasmic dialysis and the loss of key components of the intracellular signaling system. The aim of this study was to develop a simple theoretical model to estimate the expected effect on the IK1 inward-rectifying potassium current in an experiment with intact melatonin signaling. The modeling was performed using a well-established model of rat cardiomyocyte electrophysiology (Pandit et al., 2001). The maximum conductance of IK1 (gK1) channels was chosen as the target for modulation, consistent with the established mechanism of direct receptor-mediated increase in potassium conductance under the action of melatonin.Realistic modulation values were used for the modeling. The -50% value for the antagonist effect of 1 μM luzindole was obtained by direct calculation from our experimental data. The +20% value for the agonist effect (melatonin) was determined by generalizing literature data and reflects the typical expected strength of signaling pathway modulation, rather than being strictly tied to a specific concentration.It was shown that modulation of gK1 in the specified ranges leads to significant changes in IK1 amplitude in the physiologically important range of resting potentials. The developed model serves as a “computational benchmark” for validating experimental protocols, allowing one to distinguish methodological artifacts from a true lack of effect.

## Introduction

The electrical stability of the myocardium critically depends on the coordinated functioning of ion channels, among which inward-rectifying potassium channels (responsible for the IK1 current) occupy a special place [1,2]. IK1 ensures the maintenance of a stable resting potential of cardiomyocytes and protects them from spontaneous depolarization [3].

In recent years, there has been growing interest in the role of melatonin (a pineal gland hormone) in the regulation of cardiac electrophysiology and the implementation of cardioprotective effects [4, 5]. The main effect of melatonin on cells is mediated through membrane receptors MT1 and MT2, coupled with inhibitory G protein (Gi) [6]. Activation of this pathway leads to suppression of adenylate cyclase activity, a decrease in the level of intracellular cyclic AMP and modulation of protein kinase A (PKA) activity [7]. This conserved mechanism underlies the antiadrenergic and cardioprotective effects of melatonin [7].

An important consequence of melatonin receptor activation is modulation of ion channels. Studies on neurons of the suprachiasmatic nucleus (the central clock of the brain) have shown that melatonin causes a direct and rapid increase in potassium conductivity, leading to the generation of an outward current and membrane hyperpolarization [8]. This effect is receptor-mediated and is likely associated with Gi-dependent inhibition of cAMP-dependent processes [8, 9].

In experiments on isolated cardiomyocytes using the patch-clamp method in a whole-cell configuration, it is often impossible to reproduce the expected effect of melatonin on the IK1 current [10]. In our opinion, one of the key problems lies in the dialysis of the cytoplasm with the contents of the pipette, which leads to the leaching or dilution of key components of the intracellular signaling cascade (G proteins, second messengers, kinases) necessary for signal transmission from the receptor to the channel [11]. Thus, a negative result in such an experiment may be a methodological artifact rather than evidence of the absence of an effect in vivo.

The direct justification for this modeling was provided by experimental data indicating the ability of melatonin to modulate the function of cardiomyocyte ion channels. In particular, studies conducted at the Institute of Physics of the Komi Scientific Center of the Ural Branch of the Russian Academy of Sciences demonstrated that blockade of melatonin receptors with the antagonist luzindole leads to a decrease in IK1 current and a slowing of conduction in the ventricular myocardium [10]. These data suggest that melatonin itself, being an agonist of these receptors, has an opposite, enhancing effect on IK1. Thus, this modeling is based on the hypothesis that melatonin and luzindole have a direct modulating effect on the conductivity of inward rectifier channels.

The aim of this study was to develop a theoretical model to estimate the expected effect on IK1 current in an experiment if the intracellular melatonin signaling pathway remained intact. To this end, the modulation of the maximum conductance of IK1 channels (gK1) was simulated using the generally accepted mathematical model of cardiomyocytes by Pandit et al. [12] in ranges supported by literature and our own experimental data. The developed model serves as a tool for planning experiments and interpreting their results, allowing us to separate the limitations of the methodology from the true physiological mechanisms.

## Research methods

### Registration using patch clamp in the “whole cell” configuration

The experimental protocol for cell isolation, solution compositions, and IK1 current recording parameters are described in detail in our previous work. [13] To assess the acute effect of melatonin on ionic currents, nanomolar and micromolar concentrations of melatonin or 1 μM luzindole were added to each of the extracellular solutions.

### Current modeling

The IK1 current was modeled using a mathematical model of adult rat ventricular cardiomyocyte electrophysiology developed by Pandit et al. [12]. The original equation for the IK1 current was accurately reproduced according to the data provided in the Appendix of the original publication. The model was implemented in Python 3.x using the NumPy and Matplotlib libraries. The maximum conductance of gK1 was chosen as the target for pharmacological modulation. This choice is justified by the following provisions. First, in the model equation, gK1 is a scalar multiplier directly determining the current amplitude, which corresponds to the classical approach within the Hodgkin-Huxley models, where a change in this parameter reflects a change in the density of functional channels in the membrane or their unitary conductance [12]. Second, in neurons of the suprachiasmatic nucleus, the direct action of melatonin is to increase potassium conductance [8], which makes gK1 modulation a logical extrapolation of this mechanism to cardiomyocytes. Third, the melatonin signaling pathway (MT1/2 → Gi → decreased cAMP → decreased PKA activity) can modulate the phosphorylation state of Kir2.x potassium channels responsible for IK1 [1], and the final integral result of such modulation is a change in the effective maximum conductance of the channel ensemble.

To ensure quantitative agreement between the model and experimental data, the IK1 equation parameters were calibrated using the weighted sum-of-squares deviations method. The calibrated parameters were the maximum conductance gK1 and a scaling factor accounting for systematic deviations. Calibration using data obtained in the control and under the influence of the antagonist luzindole (1 μM) confirmed that the effect of luzindole is equivalent to a 50% decrease in the effective conductance gK1, which is in quantitative agreement with direct experimental measurements. The calibrated model was used to generate smooth current-voltage characteristics (CVC) over the entire range of membrane potentials (–120 to +60 mV).

In the work of Durkina et al.[10], luzindole caused a depolarization of the resting potential from –73 to –63 mV (ΔV = +10 mV). To estimate the corresponding decrease in potassium conductance, a relationship following from a simplified form of the Goldman–Hodgkin–Katz equation for the resting potential was used. Substituting ΔV = +10 mV gives an estimate of the decrease in conductance to 1/1.475≈0.68 (rounded to 0.7) of the control value. However, the simultaneously observed twofold increase in the duration of the action potential (from 136 to 288 ms) indicates a more pronounced suppression of the IK1 current, responsible for the final phase of repolarization. Therefore, a coefficient of 0.5 was adopted in the model, which is quantitatively consistent with the observed slowing of repolarization and is an estimate reflecting both experimental effects.

To simulate complete blockade of the tonic influence of endogenous melatonin, a rounded value was used. For the melatonin agonist, the change in conductance was selected based on a comprehensive assessment of the literature data. In neurons of the suprachiasmatic nucleus, melatonin (10 μM) increased potassium conductance by approximately 20% [8]. In isolated hearts, melatonin (50 μM) inhibited the isoproterenol-induced increase in cAMP by 34% [7], demonstrating the ability to significantly modulate intracellular signaling in cardiomyocytes. Given the direction of action, a modulation value of +20% (coefficient 1.2) was adopted for melatonin as a conservative estimate of its activating effect. Although direct experimental confirmation of gK1 modulation via melatonin receptors in cardiomyocytes is limited, preliminary data from our laboratory indicate the sensitivity of IK1 to this signaling pathway. Thus, the choice of gK1 as a target is a working hypothesis based on neuronal analogies [8] and our own observations.

The baseline gK1 value was set to 0.024 μS according to the Pandit model [12]. The simulations were performed in the membrane potential range from –120 to +60 mV. The current-voltage characteristics were calculated and compared for three conditions: control (gK1), melatonin treatment (gK1 × 1.2), and luzindole treatment (gK1 × 0.5).

## Research results

Modeling confirmed the characteristic IK1 current-voltage characteristic with pronounced inward rectification. The maximum current amplitude was observed in the region of negative potentials, progressively decreasing as the membrane depolarized (Fig. 1). The zero current crossing occurred near the calculated potassium equilibrium potential, consistent with the expected properties of Kir2.x channels.

**Fig. 1.**
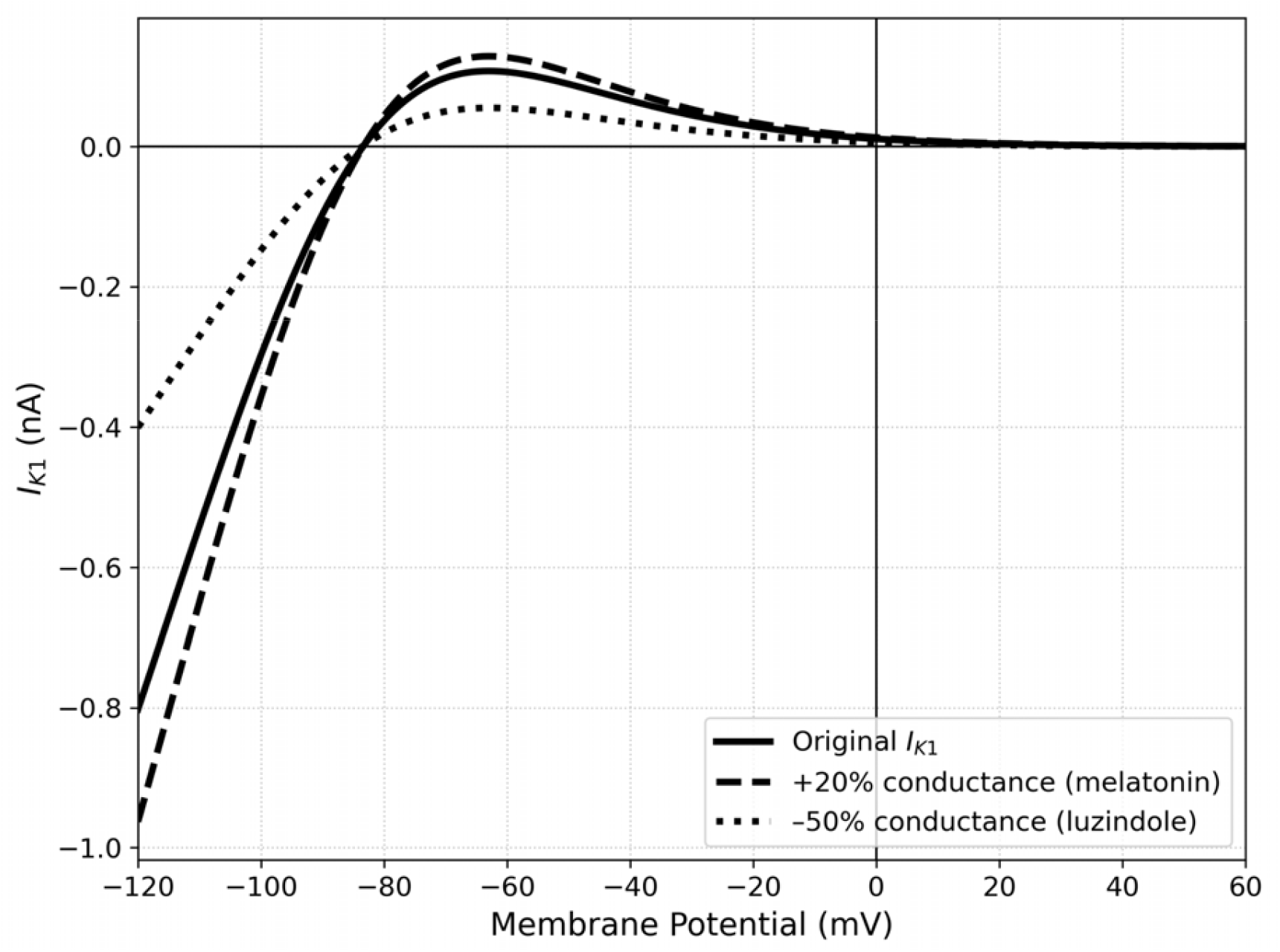
Volt-ampere dependences of IK1 current under normal conditions and with modulation of the maximum gK1 conductance. Solid line – control, dashed line – melatonin effect, dotted line – luzindole effect.

Modulation of the maximum gK1 conductance resulted in linear, proportional changes in current amplitude across the entire studied range of membrane potentials. The most significant changes in absolute current magnitude were observed in the region of physiologically significant negative potentials. When clamping the potential at –100 mV, the following quantitative results were obtained: the basal current density under control conditions was –0.299 nA. Simulated channel activation (a 20% increase in gK1) resulted in an increase in current to –0.359 nA, which exactly corresponds to the specified increment. Inhibition of conductance by 50% caused a reduction in current to –0.149 nA, which also exactly corresponded to the target modulation. A similar linear dependence was maintained at a potential of –80 mV. In the range of potentials more positive than –40 mV, where the IK1 current is virtually absent, absolute changes in amplitude were minimal, which is consistent with the inward rectification mechanism. Numerical calibration of the model using experimental data for control conditions and luzindole (1 μM) confirmed its validity (Fig. 2). Optimization yielded a gK1 value of 0.0240 μS for the control and 0.0120 μS for the luzindole condition. Thus, the effect of 1 μM luzindole is quantitatively equivalent to a 50% decrease in the effective conductance of IK1 channels, fully confirming the validity of the modulation level chosen in the model for the antagonistic effect. The limited direct experimental data on the effect of melatonin on IK1 in cardiomyocytes emphasizes the predictive value of the presented approach. The modeling results form a specific quantitative hypothesis: activation of the melatonin signaling pathway can lead to an increase in gK1 conductance by approximately 20%. The resulting simulated I–V characteristics and calculated current amplitude values serve as a benchmark for future experimental testing. The proposed model is a working tool that allows us to translate qualitative understanding of ion channel modulation through intracellular signaling cascades into quantitative predictions of functional electrophysiological effects at the cellular level.

**Fig. 2.**
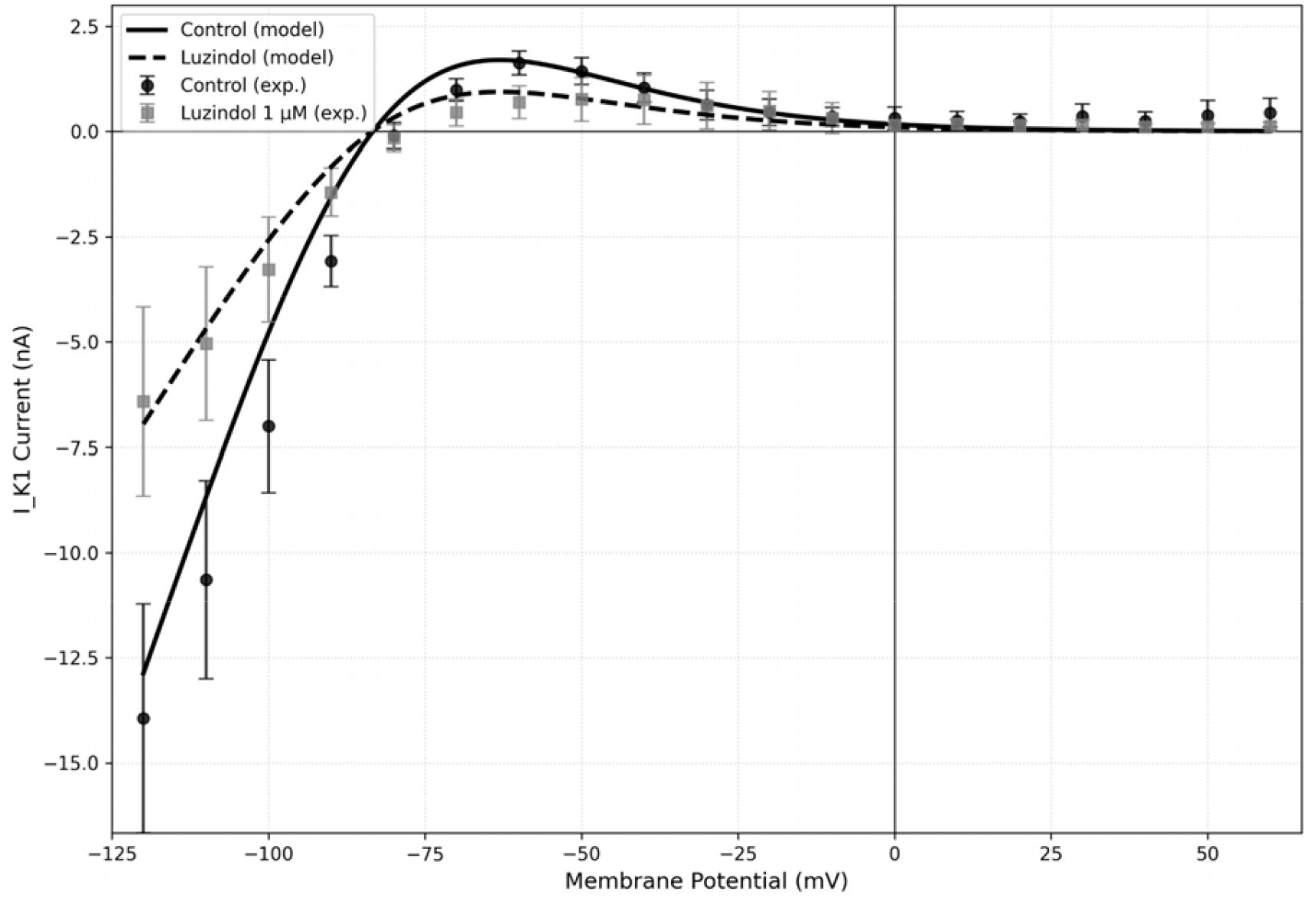
Calibration of the IK1 current model using experimental data. The graph shows the experimental current-voltage characteristics (mean values ± standard deviation) for the control and 1 μM luzindole conditions (black circles). The lines correspond to the fitted model curves after numerical optimization of the parameters. Calibration confirms that the effect of luzindole quantitatively corresponds to a 50% reduction in effective conductance.

## Discussion of results

The main result of this study is the development of a theoretical standard for predicting the effect of melatonin and luzindole on the IK1 current under conditions of preserved intracellular signaling. The presented model demonstrates that modulation of the maximum conductance of IK1 channels (gK1) in the range of +20% for the agonist and -50% for the antagonist leads to statistically significant changes in the current amplitude in the physiologically critical range of resting potentials (from -100 to -80 mV). The increase in IK1 under the influence of melatonin is consistent with its known cardioprotective properties, which are likely mediated by stabilization of the resting potential and suppression of spontaneous electrical activity [4]. Conversely, a decrease in IK1 by luzindole creates an electrophysiological substrate for destabilization of the membrane potential, which can be considered a potentially proarrhythmic factor.

The developed model has significant methodological significance. The data obtained are essential for the correct planning and interpretation of patch-clamp experiments. A known limitation of the whole-cell configuration is cytoplasmic dialysis, which leads to the loss of low-molecular components of signaling cascades (G proteins, cAMP, ATP) [1], which can cause false-negative results when studying receptor-mediated effects. This model provides a specific quantitative prediction that can be used to assess the adequacy of the experimental protocol. If the observed effect of melatonin or luzindole matches the model prediction in direction and order of magnitude, this indicates the integrity of the intracellular signaling pathway under the conditions of a particular experiment. If the effect is absent or significantly weaker than predicted, this serves as a strong argument in favor of the methodological limitations associated with dialysis and indicates the need to modify the protocol (e.g., switching to a perforated patch configuration). It should be especially emphasized that the value of gK1 modulation under the action of the agonist chosen for the modeling (+20%) is not a prediction for a strictly defined concentration of melatonin, but a reasonable quantitative benchmark for the strength of the effect expected from the activation of this signaling pathway in principle. This estimate is based on extrapolation of data from different experimental models: in neurons, 10 μM melatonin caused an increase in potassium conductivity by approximately 20% [8], and in cardiomyocytes, 50 μM melatonin demonstrated the ability to significantly inhibit cAMP-dependent processes [7]. Thus, the model predicts not the effect of a specific dose, but rather the order of magnitude of change in current amplitude (tens of percent) that should be considered a physiologically plausible marker of successful signal transmission from the receptor to the channel under intact-cell conditions. This is a key aspect for the correct use of the model as a validation tool: if, in an experiment with preserved signaling, the observed effect of melatonin is comparable in direction and order of magnitude to the reference (+20%), the protocol can be considered adequate.

The proposed approach also helps explain a frequently observed experimental phenomenon: the presence of a pronounced antagonist effect of luzindole in the absence of the agonist effect of melatonin. This may be due to the fundamentally different dependence of their action on the preservation of the intracellular environment. As a competitive antagonist, luzindole exerts its effect primarily through direct binding to the receptor, blocking its tonic activity. This initial stage can occur even with partial leakage of intracellular components. In contrast, the implementation of the melatonin agonist effect requires not only binding to the receptor, but also subsequent full activation of the entire intracellular cascade (G protein → adenylate cyclase → cAMP → PKA → target channel), each link of which is vulnerable to dialysis in the whole-cell configuration [1]. Thus, the model emphasizes that the absence of the melatonin effect in the presence of the luzindole effect does not refute the existence of the signaling pathway*in vivo*, but, on the contrary, indicates its sensitivity to methodological artifact.

The obtained quantitative agreement between the predicted indole loss effect by the model (–50%) and the experimental data (–53%) [12] confirms the adequacy of the chosen approach—modeling through modulation of the scalar parameter gK1. This indicates that the experimentally observed decrease in IK1 upon receptor blockade is primarily due to a decrease in the maximum channel conductance. The slight discrepancy may be due to additional effects not accounted for in the model or to nonlinearity of signaling. Importantly, the model parameterized based on antagonist data provides a plausible quantitative estimate of the agonist effect, which requires direct experimental verification under conditions of a preserved intracellular signaling pathway.

The developed model is simplified and has several limitations. It does not take into account possible non-receptor (e.g., antioxidant) effects of melatonin. The effect on gK1 is modeled as instantaneous and constant, whereas*in vivo*Temporal dynamics and desensitization may be observed. The model focuses exclusively on IK1, while melatonin can modulate other ionic currents, exerting complex effects on cardiomyocyte electrophysiology. A promising direction is the integration of this approach into full-fledged action potential models to assess the integral effects on action potential duration and rhythm stability, as well as further experimental studies to accurately quantify the contribution of the Gi/cAMP-dependent pathway to IK1 modulation.

It is important to emphasize that the developed model is purposefully predictive and methodological in nature. Its primary objective is not to retroactively describe the full array of experimental data already obtained, but to propose a quantitative computational benchmark for planning and interpreting experiments aimed at studying receptor-mediated modulation of ion channels. The availability of preliminary experimental data qualitatively and quantitatively consistent with the model’s predictions, including experiments modulating the cAMP-dependent pathway (e.g., using forskolin), confirms the validity of the chosen approach. However, these data have been deliberately omitted from this publication to avoid duplication and maintain the article’s clear focus on the development, validation, and verification of the modeling method as an independent tool. A full experimental verification of all the model’s implications and its integration into more complex descriptions of cardiomyocyte electrophysiology are the subject of a separate, large-scale study, the results of which will be presented in future papers. Thus, this article formulates a rigorous quantitative hypothesis and provides researchers with a tool for testing it under conditions that minimize methodological artifacts.

## Conclusion

The main result of this study is the development of a theoretical model that quantitatively describes the modulation of the maximum conductance of IK1 (gK1) channels under the influence of a melatonin receptor agonist (melatonin, +20%) and antagonist (luzindole, -50%). The model, implemented based on the generally accepted formalism of Pandit et al. (2001), was directly numerically calibrated using experimental data for luzindole, confirming the adequacy of the chosen approach. The developed model serves as a predictive computational benchmark for the patch-clamp methodology. Its practical value lies in providing a specific quantitative benchmark for differentiating methodological artifacts associated with whole-cell cytoplasmic dialysis from the true absence of a receptor-mediated effect. The availability of preliminary experimental data consistent with the model’s prediction confirms the validity of the hypothesis; however, a detailed presentation of these data is beyond the scope of this methodological work and would require a separate experimental study. Thus, the model provides a tool for planning and validating experiments aimed at studying the intracellular signaling regulation of ion channels in cardiomyocytes.

## Authors’ contributions

The idea and planning of the article (M.A.G., V.F.S.), conducting experiments and processing data (M.A.G.), modeling and preparation of illustrations (V.F.S.), preparation and editing of the manuscript (M.A.G., V.F.S.).

## Data availability

The model source code and experimental data are available in the GitHub repository:https://github.com/victoriasafiulina-design/VictoriaSafiulina/blob/main/IK1.py https://github.com/victoriasafiulina-design/VictoriaSafiulina/blob/main/Luz%20(2).py

